# Development of amidase-dependent pyrazinoic acid prodrugs with activity against pyrazinamide resistant *Mycobacterium tuberculosis*

**DOI:** 10.1101/2022.07.05.496887

**Authors:** Carly Levine, Ravindra Jadhav, Yan Pan, Kholiswa Tsotetsi, Xin Wang, Divya Awasthi, Courtney Grady, Anil Shelke, Samer S. Daher, Todd Richmann, Riju Shrestha, Paridhi Sukheja, Jimmy Patel, Pamela R. Barnett, Ryan J. Dikdan, Thomas Kim, Riccardo Russo, Jennifer Hanna, Matthew Zimmerman, Véronique Dartois, Joel S. Freundlich, David Alland, Pradeep Kumar

## Abstract

Rapid emergence of drug resistance in *Mycobacterium tuberculosis* (Mtb) is one of the most significant healthcare challenges of our time. The cause of drug resistance is multifactorial, with the long course anti-tubercular therapy required to treat tuberculosis (TB) constituting a major contributing factor. Introduction of pyrazinamide (PZA) resulted in shortening of TB treatment from twelve to six months and consequently played a critical role in curbing drug resistance that developed over long course therapy. Nevertheless, because PZA is a prodrug activated by a nonessential amidase, PncA, resistance to PZA develops and frequently results in treatment failure. Here, we leveraged a whole cell drug screening approach to identify anti-tuberculars with unconventional mechanisms of action or activation that could be further developed into compounds effective at killing Mtb resistant to PZA. We discovered an amide containing prodrug, DG160, that was activated by the amidase, Rv2888c (AmiC). This amidase was capable of metabolizing a variety of amide containing compounds including a novel pyrazinoic acid-isoquinolin-1-amine prodrug, JSF-4302, which we developed as a potential PncA-independent replacement for PZA. As predicted, AmiC activation of JSF-4302 led to the generation of POA in Mtb including in a PZA resistant clinical isolate, thereby successfully delivering the active component of PZA while bypassing the need for activation by PncA. This work provides a framework for a new approach to drug development and prodrug activation in Mtb.

**SIGNIFICANCE:** Pyrazinamide (PZA) is a vital component of *Mycobacterium tuberculosis* (Mtb) treatment since its inclusion shortened tuberculosis therapy by six months. However, PZA is a prodrug and resistance develops at a high frequency due to mutations in its activator PncA. Here, we present the discovery of amide-containing anti-tubercular prodrugs that are activated intracellularly by the Mtb amidase, AmiC. Taking advantage of this finding, we successfully designed and synthesized pyrazinoic acid (POA) prodrugs that were activated by AmiC and found that these compounds delivered intracellular POA to PZA-resistant Mtb isolates that contained a nonfunctional PncA. This new approach to prodrug development provides a method for delivering conjugated drugs into Mtb with the potential to overcome clinical drug resistance.

## INTRODUCTION

Tuberculosis (TB) is responsible for roughly 1.5 million deaths annually, second only to SARS-CoV-2 as the leading cause of death due to an infectious agent (1). However, TB has constituted a pandemic for more than two decades. Firstline treatment against *Mycobacterium tuberculosis* (Mtb), the etiological agent of TB, is a six-month multidrug regimen consisting of isoniazid (INH), ethambutol (EMB), rifampicin (RIF), and pyrazinamide (PZA). Recent experimental evidence suggests that treatment duration can be shortened from six to four months via the substitution of RIF with high dose rifapentine and EMB with moxifloxacin (2); however, PZA remains an essential component of this shortened regimen. The global emergence of multidrug-resistant TB (MDR-TB) that is resistant to both INH and RIF (3), extensively drug-resistant (XDR), and totally drug-resistant TB threaten the long term success of any new treatment (4–7). Thus, there is a crucial need to develop novel anti-tubercular drugs that are active against drug resistant Mtb and can further shorten treatment duration.

Many of the compounds currently in use to treat TB are prodrugs that require activation by Mtb enzymes. These include INH, PZA, ethionamide and pretomanid (8–10). PZA is especially important because its introduction enabled the shortening of TB therapy from twelve to six months, likely due to its ability to target persistent Mtb populations (11–13). PZA is converted within Mtb by the bacterial pyrazinamidase (PncA) into its active form pyrazinoic acid (POA), whose mechanism of action may be characterized by a complex polypharmacology (14–17). The main driver of clinical resistance to PZA is through development of mutations in the PncA encoding gene, *pncA* (18–22). Resistant mutants raised on high concentrations of POA *in vitro*, as well as roughly 10% of PZA-resistant clinical Mtb isolates, can also develop mutations in *panD* (23–25). The *panD* gene encodes for an aspartate decarboxylase which carries out the penultimate step in pantothenate biosynthesis, an essential precursor of coenzyme A synthesis. Resistance to PZA develops in about 2.5% of fully susceptible TB cases and increases to 60% when treating MDR-TB (26). Therefore, an urgent need exists to design improved POA delivery systems that can be activated by alternative enzymes.

Contemplating various modes of delivery of POA, we were drawn to published studies of the hydrolysis of ester-, amide-, or carbamate-containing prodrugs into their active components in many species ranging from humans to bacteria (27–29). These types of prodrugs typically have favorable pharmacokinetic properties and were designed to increase cell permeability by masking polar groups (30–32). The Mtb genome encodes for over 150 putative hydrolases with predicted protease, amidase, or esterase function based on Pfam and experimental evidence (33). In other bacteria, many hydrolases are multifunctional and can nonspecifically metabolize a broad range of substrates (34, 35). However, this type of substrate promiscuity is not yet well documented in Mtb.

Here, we identified an amide containing compound DG160 that was active against Mtb with a minimum inhibitory concentration (MIC; the lowest compound concentration to achieve 90% reduction in bacterial growth in culture) in the low micromolar range. Further evaluation of drug-resistant mutants revealed that DG160 was a prodrug activated by the amidase Rv2888c, AmiC which has also been shown to metabolize indole-4-carboxamide prodrugs (36). Here we show that AmiC is able to activate or deactivate a variety of amide containing compounds; and we exploit AmiC to develop novel prodrug hybrids of the DG160 amine bound to POA. These efforts produced a new synthetic POA prodrug, JSF-4302, which successfully released intracellular POA in Mtb and showed potent *in vitro* activity against a PncA-deficient, PZA resistant, clinical Mtb isolate.

## RESULTS

### AmiC is a novel prodrug activator

We screened a GlaskoSmithKline (GSK) library of anti-tubercular compounds (37) with the *Mycobacterium bovis* BCG (BCG) P*iniBAC* reporter strain to identify compounds that targeted Mtb cell wall biosynthesis (38). This work highlighted a moderate inducer of P*iniBAC*, SB-746177, which we renamed DG160 (Fig. S1). DG160 had a potent MIC of 0.48 μg/mL (1.6 μM) against wild type (WT) Mtb strain H37Rv as well as most clinical Mtb isolates (Table 1 and Table S1), as measured using an alamarBlue assay. DG160 was bactericidal, causing a 2 log_10_ reduction in colony forming units (CFU) after 4 days of treatment at 10x MIC (Fig. S2). Mutants resistant to DG160 occurred at a frequency of ∼1x10^-7^ when plated on DG160 at 16x the MIC, and all harbored mutations in *amiC* (*rv2888c*) which encoded a putative amidase. Interestingly, 4 clinical Mtb strains that were resistant to DG160 also harbored mutations in *amiC* (Table S1).

**Table 1:** Efficacy, genotype, and cross-resistance of selected amide containing compounds in various Mtb strains.

| Strain | | | | alarmarBlue MIC (μg/mL) | | | |
| --- | --- | --- | --- | --- | --- | --- | --- |
|  | Type | Gene Mutated | Amino Acid Change | INH | DG160 | DA60 | DG85 |
| Mtb H37Rv | Laboratory | NA | NA | 0.04 | 0.48 | 1.05 | 0.66 |
| Mtb H37Rv:pVV16 | Laboratory | NA | NA | 0.04 | 0.96 | 1.05 | 0.66 |
| Mtb H37Rv:pVV16-*amiC* | Laboratory | NA | NA | 0.04 | 0.12 | 0.26 | 2.7 |
| Mtb H37Rv DRM-DG160.4 | Laboratory | *amiC*<sup>+</sup> | H346fs | 0.04 | 7.6 | 8.4 | 0.33 |
| Mtb H37Rv DRM-DG160.8 | Laboratory | *amiC* | L402P | 0.04 | 7.6 | ND | ND |
| Mtb H37Rv DRM-DG160.9 | Laboratory | *amiC* | K84fs | 0.04 | 7.6 | ND | ND |
| Mtb H37Rv DRM-DG160.12 | Laboratory | *amiC* | V417fs | 0.04 | 7.6 | ND | ND |
| Mtb DRM-PZA | Clinical | *pncA*<sup>†</sup> | Y34fs | >2.8 | 0.48 | ND | ND |
| Mtb CDC1551 | Laboratory | NA | NA | 0.04 | 0.96 | 4.2 | ND |
| Mtb CDC1551-*amiC*::tn | Laboratory | *amiC* | A260fs | 0.04 | 7.6 | 8.4 | ND |

\* Other mutations detected
† Whole genome sequenced
ND: not done
NA: not applicable

DG160 contains a central amide linker that we hypothesized was susceptible to amidohydrolysis by AmiC which would then result in the formation of fusaric acid and isoquinolin-1-amine (Fig. 1A). Analysis of the intrabacterial drug metabolism (IBDM) (39, 40) of DG160 via liquid chromatography tandem mass spectrometry (LC-MS/MS) showed that WT Mtb rapidly hydrolyzed DG160 in a time-dependent manner and that the proposed metabolites were indeed formed (Fig. 1B). A DG160 drug-resistant mutant that contained an *amiC*-H346 frameshift (fs) mutation (DRM-DG160.4) metabolized DG160 significantly slower than WT Mtb (Fig. 1C). However, metabolite levels in both strains were comparable at the final 24 h time point (Fig. 1B and 1C). These data indicate that although AmiC is the primary amidohydrolase acting on DG160, Mtb likely expresses additional amidases that are also capable of hydrolyzing DG160. Conversely, an *amiC* overexpressing strain (H37Rv:pVV16-*amiC*) showed faster amide hydrolysis compared to its vector control (H37Rv:pVV16) (Fig. 1E and 1D). These data were also quantified by calculating the area under the curve (AUC0-24h) of DG160 over the experimental time course. The results showed significantly slower metabolism of DG160 by DRM-DG160.4 compared to WT Mtb, and faster metabolism by H37Rv:pVV16-*amiC* (Fig. 1F). It should be noted that DG160 was stable in cell free media over 24 h. The MIC of the produced metabolites, fusaric acid and isoquinolin-1-amine, in WT Mtb were 36 µg/mL (200 µM) and 14.4 µg/mL (100 µM), respectively, suggesting that prodrug activation within the bacteria was essential for the activity of DG160.

**Fig. 1:**
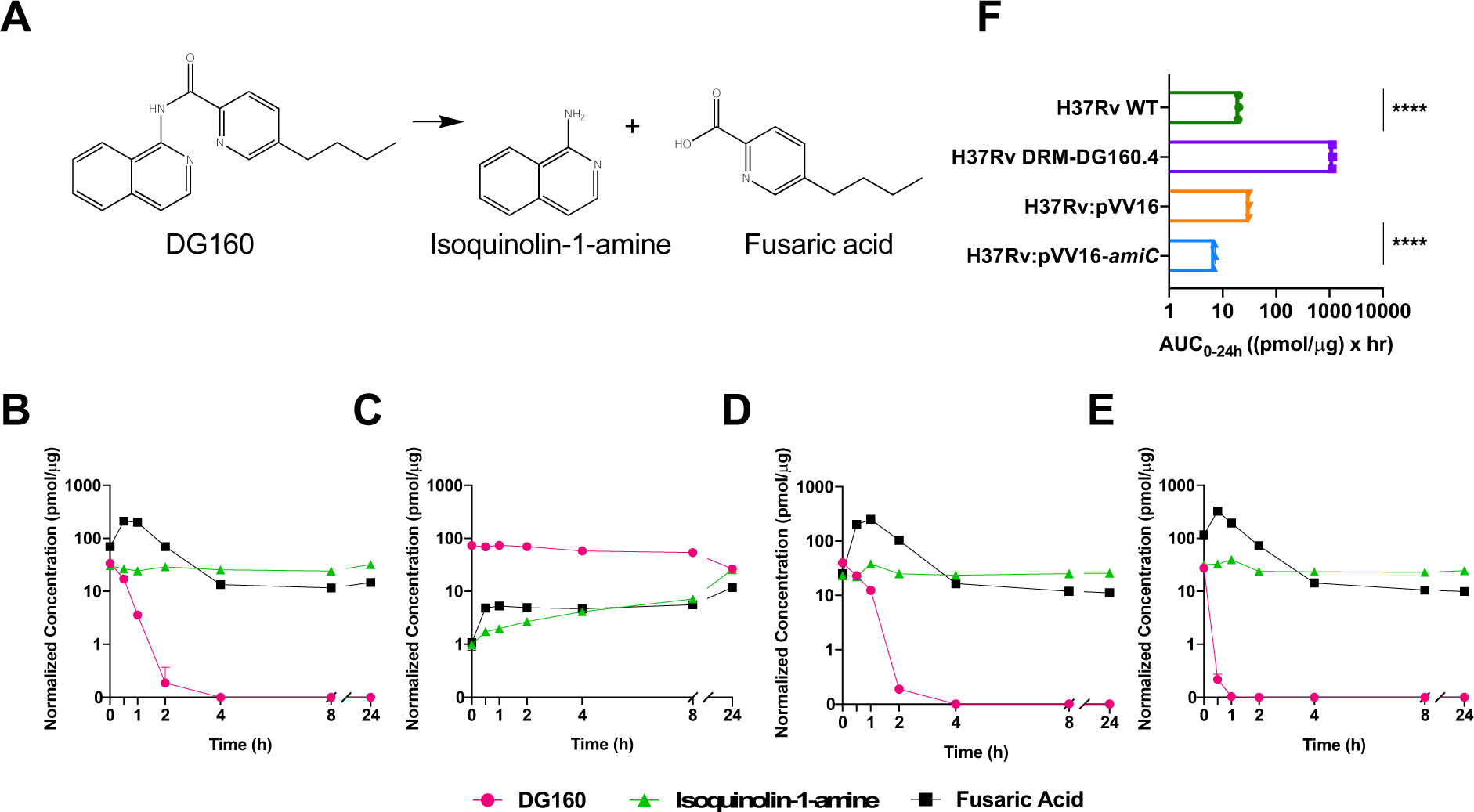
Intrabacterial amidohydrolysis of DG160 by AmiC. **A)** Schematic of amide hydrolysis and subsequent generation of isoquinolin-1-amine and fusaric acid. **B)** Mtb H37Rv WT, **C)** Mtb H37Rv DRM-DG160.4, **D)** Mtb H37Rv:pVV16, **E)** Mtb H37Rv:pVV16- *amiC,* **F)** AUC_0-24h_ of DG160 over the 24 h time course. Pink circles: DG160, green triangles: isoquinolin-1-amine, black squares: fusaric acid. Data points and error bars represent the mean and standard deviation, respectively. At all timepoints n = 3. Two-way ANOVA was used for statistical analysis. **** p<0.0001.

### Purified AmiC hydrolyzed DG160 and other diverse amide bond containing compounds

Several compounds in our collection of active anti-tuberculars contain amide bonds and thus, we wondered if AmiC could indiscriminately activate these other molecules. Testing MIC, we found that H37Rv:pVV16-*amiC* was 4x more susceptible to SMR000323973 (41, 42) (renamed DA60) and 4x more resistant to GSK426032A (37) (renamed DG85) compared to WT Mtb (Table 1). We next tested whether AmiC could hydrolyze DG160, DA60, and/or DG85 *in vitro*. The C-terminus His6-tagged AmiC was purified from H37Rv:pVV16-*amiC* (Fig. S3). AmiC hydrolyzed DG160 into the expected fusaric acid and isoquinolin-1-amine metabolites (Fig. S4). AmiC also fully metabolized DA60 after a 2 h incubation (Fig. S5) and only partially metabolized DG85 (Fig. S6). These data confirmed that AmiC could metabolize a spectrum of amide containing small molecules, likely activating the anti-tubercular properties of DG160 and DA60 and inactivating the anti-tubercular activity of DG85.

### Design, synthesis, and characterization of JSF-4302

The substrate promiscuity of AmiC and potential existence of other amidases in Mtb including at least three AmiC homologs (Fig. S7) suggested a pathway to design novel anti-tubercular prodrugs that could release an active moiety after hydrolysis by AmiC. As a test of this prodrug design scheme, we designed a hybrid compound that could deliver the active acid component of PZA, POA, into Mtb independently of PncA hydrolysis, with the intention of reviving the activity of PZA in *pncA* mutated, PZA-resistant Mtb strains. We designed and synthesized a series of POA amides. A subset of them (JSF-4301 – JSF-4304; Table 2 and Table S2) featured the isoquinoline-1-amine of DG160 (JSF-4302) or a 2-aminopyridine derivative (JSF-4301, 4303, 4304).

**Table 2:**
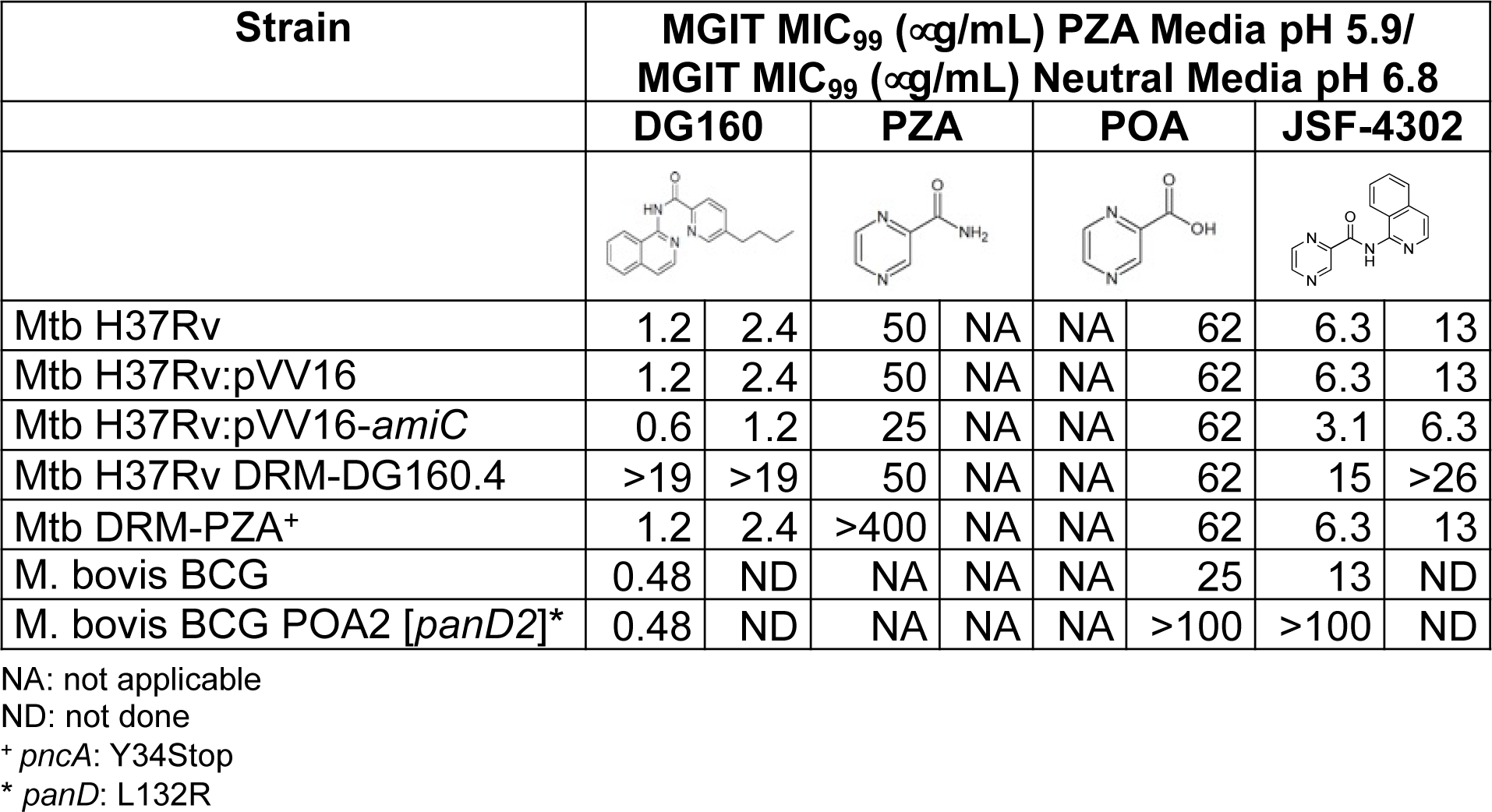
Efficacy of POA derivatives using BACTEC MGIT.

| Strain | MGIT MIC<sub>99</sub> (μg/mL) PZA Media pH 5.9/<br>MGIT MIC<sub>99</sub> (μg/mL) Neutral Media pH 6.8 | | | | | | | |
| --- | --- | --- | --- | --- | --- | --- | --- | --- |
|  | DG160 |  | PZA |  | POA |  | JSF-4302 |  |
| Mtb H37Rv | 1.2 | 2.4 | 50 | NA | NA | 62 | 6.3 | 13 |
| Mtb H37Rv:pVV16 | 1.2 | 2.4 | 50 | NA | NA | 62 | 6.3 | 13 |
| Mtb H37Rv:pVV16-*amiC* | 0.6 | 1.2 | 25 | NA | NA | 62 | 3.1 | 6.3 |
| Mtb H37Rv DRM-DG160.4 | >19 | >19 | 50 | NA | NA | 62 | 15 | >26 |
| Mtb DRM-PZA<sup>+</sup> | 1.2 | 2.4 | >400 | NA | NA | 62 | 6.3 | 13 |
| M. bovis BCG | 0.48 | ND | NA | NA | NA | 25 | 13 | ND |
| M. bovis BCG POA2 [*panD2*]<sup>*</sup> | 0.48 | ND | NA | NA | NA | >100 | >100 | ND |

NA: not applicable
ND: not done
<sup>+</sup> *pncA*: Y34Stop
<sup>\*</sup> *panD*: L132R

We found that JSF-4301, JSF-4302, JSF-4303, and JSF-4304 had moderate activity with MIC values of 160 μg/mL (800 µM), 25 μg/mL (100 µM), 172 μg/mL (800 µM), 43 μg/mL (200 µM), respectively, when tested using the alamarBlue assay (Table S2). However, PZA is only active *in vitro* at an acidic pH (43). Therefore, we tested the MIC99 (MIC needed to inhibit 99% growth of the bacterial culture) of these compounds using the mycobacterial growth indicator tube (MGIT) system with MGIT PZA Media at pH 5.9. Using this acidic media, we found that JSF-4301, JSF-4302, JSF-4303, and JSF-4304 had MIC99 values of 4.9 μg/mL (25 µM), 6.3 μg/mL (25 µM), 27 μg/mL (130 µM), and 5.4 μg/mL (25 µM), respectively (Table 2 and Table S3). Importantly, these activities were superior to PZA (MIC99 50 µg/mL (410 µM)) under the same conditions.

**Table 3:**
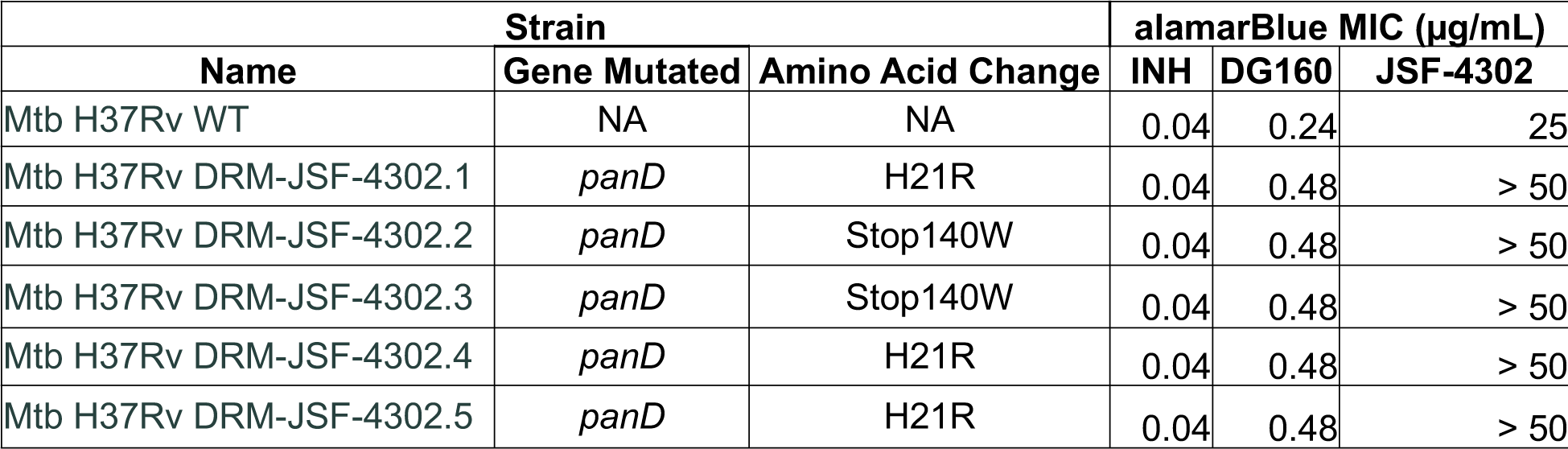
Efficacy, genotype, and cross-resistance of JSF-4302 resistant mutants

We focused on JSF-4302 for subsequent studies because it had a reasonable MIC99 and minimal cytotoxicity with a CC50 (i.e., the minimum amount of compound needed to inhibit the growth of the mammalian cell line by 50%) of 66 μg/mL versus J774 macrophages and 70 μg/mL against Vero CCL81 kidney epithelial cells, resulting in both selectivity indices exceeding 10. Consistent with its design as an AmiC-dependent prodrug, the activity of JSF-4302 was reduced in the DRM-DG160.4 strain with an MIC99 of 15 μg/mL (62 µM), and its activity was increased in the *amiC* overexpression strain with an MIC99 of 3.1 μg/mL (12 µM) (Table 2). Additionally, the MIC99 of JSF-4302 in MGIT neutral media (pH 6.8) was 2-fold higher than the MIC99 in MGIT acidified media (pH 5.9) (Table 2). These data suggest that AmiC successfully metabolizes JSF-4302 in Mtb and that its activity may be partially dependent on environmental pH.

### JSF-4302 is a POA delivering prodrug

We next determined whether JSF-4302 activity was mediated by intracellularly formed POA. BCG is naturally resistant to PZA due to a non-functional PncA, but it is fully susceptible to POA (21, 44). We postulated that JSF-4302 would retain activity in BCG but would lose activity in a BCG strain that was resistant to POA due to a mutation in the *panD* gene (*panD* L132R) (45). The MIC99 of JSF-4302 in WT BCG was 13 μg/mL (52 µM) whereas the MIC99 in the POA resistant BCG strain was greater than 100 μg/mL (400 µM) (Table 2). We next evaluated the activity of JSF-4302 in a PZA resistant Mtb clinical isolate (DRM-PZA, *pncA* Y34fs), and found that it was fully susceptible to POA (MIC99 62 μg/mL (490 µM)) as well as JSF-4302 (MIC99 6.26 μg/mL (25 µM) (Table 2).

To further determine whether JSF-4302 was a POA delivering prodrug, we studied the extent to which JSF-4302 was metabolized into isoquinolin-1-amine and POA (Fig. 2A). First, using isolated AmiC, we found that JSF-4302 was successfully metabolized by AmiC *in vitro* (Fig. S8). We next used the IBDM approach to determine the metabolism of JSF-4302 within WT, DRM-DG160.4, DRM-PZA, H37Rv:pVV16, and H37Rv:pVV16- *amiC* Mtb strains. We found that JSF-4302 was metabolized by all strains, producing both isoquinolin-1-amine and POA (Fig. 2B-F). However, the extent of metabolism of JSF-4302, as correlated with its AUC0-24h value, by DRM-DG160.4 was significantly less than by WT, while H37Rv:pVV16-*amiC* metabolized JSF-4302 to a greater extent than by the H37Rv:pVV16 control strain (Fig. 2G). Thus, our results confirm that JSF-4302 is hydrolyzed largely in an AmiC-dependent manner and that POA is released in a Mtb *pncA* mutant (DRM-PZA) which strongly suggests that JSF-4302 can function as an alternative POA delivery system.

**Fig. 2:**
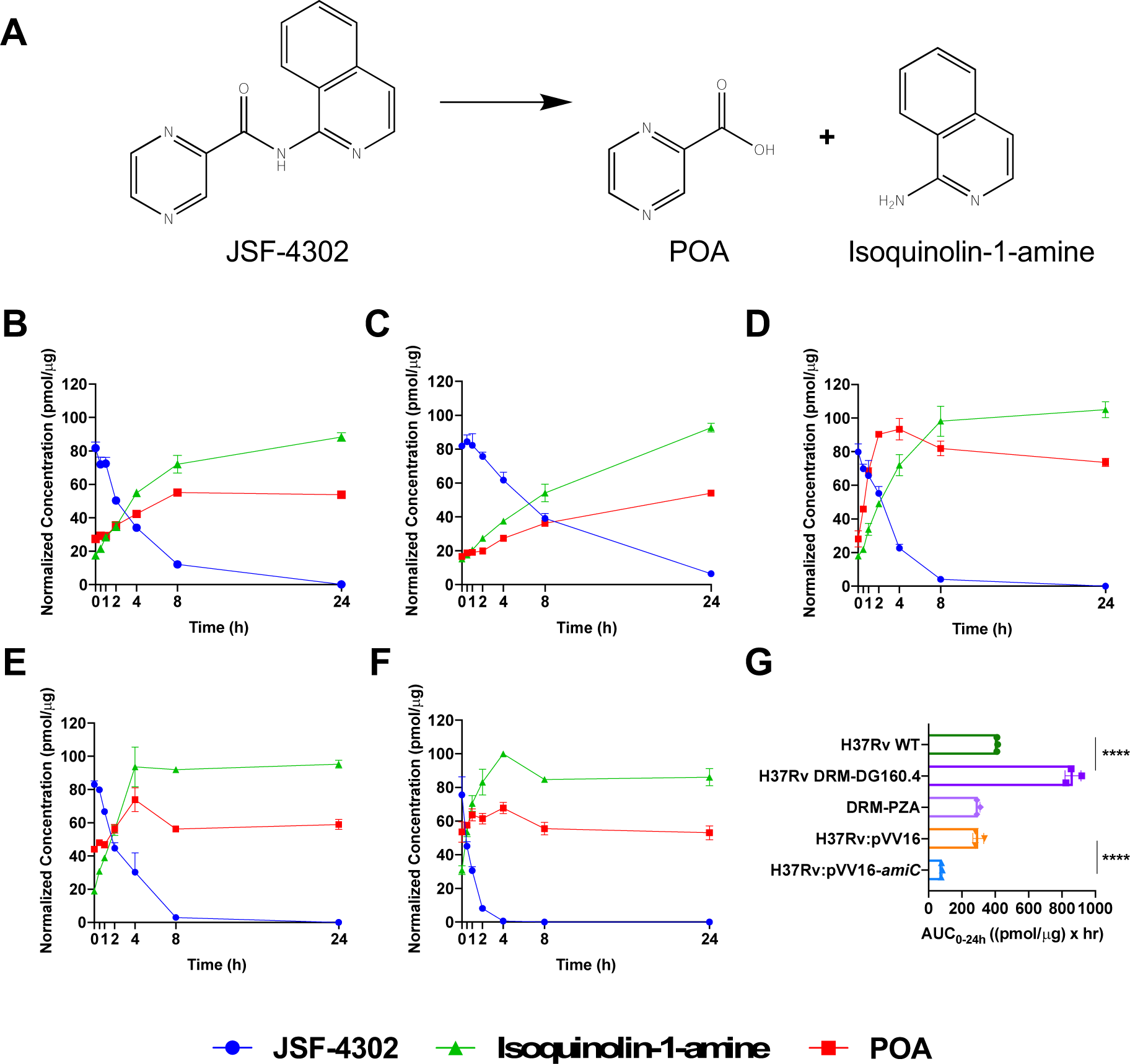
Intrabacterial metabolism of JSF-4302 and the release of POA intracellularly. **A)** Schematic of amide bond hydrolysis and subsequent generation of POA and Isoquinolin-1-amine, **B)** Mtb H37Rv WT **C)** Mtb H37Rv DRM-DG160.4, **D)** Mtb DRM-PZA, **E)** Mtb H37Rv:pVV16, **F)** Mtb H37Rv:pVV16-*amiC*, **G)** AUC_0-24h_ of JSF-4302 over the 24 h time course. Blue circles: JSF-4302, green triangles: Isoquinoline-1-amine, red squares: POA. Data points and error bars represent the mean and standard deviation, respectively. At all timepoints n = 3. Two-way ANOVA was used for statistical analysis. **** p<0.0001.

In further support of our hypothesis that JSF-4302 releases POA, Mtb was plated on 8x MIC (200 μg/mL) of JSF-4302 at a neutral pH 6.8. We found that colonies growing on these plates harbored mutations in *panD* and were greater than 2-fold resistant to the parent compound while being only low level resistant to DG160 (Table 3). Taken together, these data strongly suggest that JSF-4302 can function as an alternative POA delivery system.

## DISCUSSION

MDR-TB now accounts for almost 4% of all new cases of Mtb worldwide and has a treatment success rate of only 57% (1). Thus, the development of novel anti-tubercular compounds active against these highly drug-resistant strains is critically needed. The *amiC* gene encodes for an amidase with serine hydrolase activity (46, 47). Bioinformatics analysis of AmiC shows that it contains an amidase family signature (conserved KSS residue) found in fatty acid amide hydrolases in mammals but experimental evidence for its role in Mtb physiology is lacking (48). Here, we found that AmiC was capable of promiscuously activating or deactivating compounds containing amide bonds. Taking our findings into consideration, it is interesting to note that a recent study also attributed activation of indole-4-carboxamide prodrugs to AmiC-mediated hydrolysis (36). This group found that AmiC was able to metabolize a variety of indole-4-carboxamide derivatives that varied in size, polarity, and position of other substituents. Together these data highlight the potential importance of AmiC in the metabolism of amide containing anti-tuberculars.

First-line TB drug PZA is also activated by an amidase, PncA, which led us to propose the design of POA-isoquinolin-1-amine prodrugs that could deliver POA, the active component of PZA, but would predominantly be activated by AmiC instead of PncA. In this study, we designed a POA prodrug scaffold that successfully delivered POA to both PZA susceptible and PZA resistant strains via activation by AmiC. To further demonstrate that POA was the active moiety of JSF-4302, we raised drug-resistant mutants to JSF-4302 at a neutral pH and all harbored mutations in *panD*, the proposed target of POA. Clinically, *panD* mutants are quite rare, making up only 10% of PZA-resistant Mtb isolates (23–25).

Amide bonds are present in a number of FDA approved anti-tubercular drugs including RIF, PZA, INH, capreomycin, amikacin and β-lactams. The Mtb genome encodes for 3 homologs of AmiC along with many other amidases, and we propose that the slow but eventual breakdown of DG160 and JSF-4302 by DRM-DG160.4 may be attributed to amidohydrolysis by one of these other amidases. Overexpression of a different amidase, AmiD, caused 2-fold hyper-susceptibility to DG160 suggesting this amidase may be involved in DG160 metabolism (data not shown). Additionally, the lack of *amiC* mutations in any JSF-4302-resistant mutants combined with the confirmed release of POA highlight the potential importance of other amidases in JSF-4302 activation. Interestingly, frameshift mutations in *amiC* are associated with MDR-TB suggesting a potential role for AmiC in the metabolic activation of current anti-tubercular drugs (49). These data expose the complex capability of Mtb amidases to activate or deactivate amide containing compounds and provide a framework for developing new versions of old anti-tuberculars.

## METHODS

### Chemical and compound library *iniBAC* screen

Libraries of structurally diverse small molecules with known whole-cell anti- tubercular activity were obtained from GSK (37) and the Southern Research Institute (SRI) (41, 42). For screening, the BCG fluorescent reporter strain harboring mWASABI under the *iniBAC* promoter was cultured to an OD595 = 0.2 in Middlebrook 7H9 medium (BD Biosciences) supplemented with 10% Middlebrook oleic acid, albumin, dextrose, catalase (OADC) (BD Biosciences) and 0.05% Tween 80 (Sigma Aldrich). This culture was then dispensed into 96-well black clear-bottom half-area plates (Corning) at 90 μL/well. Compounds from the test library were then added to wells in 10 μL aliquots for a final test concentration of 10 μM. Positive and negative controls were included in each plate in triplicate. Plates were incubated for 24 h at 37 °C, and the fluorescence of each well was recorded at an excitation wavelength of 488 nm and an emission wavelength of 509 nm in a SpectraMax M5 microplate reader. Compounds were scored based on fold induction over vehicle control. The library was screened in singlets due to limitations in the amount of compound available. However, each positive hit was repeated in triplicate.

### Synthesis of small molecules

#### DG160

To a solution of isoquinolin-1-amine (36.3 mg, 0.252 mmol) in 1 mL of dichloromethane was added the hydrochloric acid salt of 1-ethyl-3-(3- dimethylaminopropyl) carbodiimide (EDC•HCl) (48.3 mg, 0.252 mmol) and 5- butylpicolinic acid (45.1 mg, 0.252 mmol). 4-Dimethylaminopyridine (DMAP) (31.0 mg, 0.252 mmol) was added to the above reaction mixture which was stirred overnight at 40 °C. After completion of the reaction as per TLC, the reaction mixture was diluted with dichloromethane, transferred to a separatory funnel, and the organic layer was washed with water, aqueous sodium bicarbonate, and saturated aqueous brine solution. The organic layer was dried over anhydrous magnesium sulfate, filtered, and concentrated *in vacuo*. The crude product was purified by flash chromatography on silica gel (0% to 70% ethyl acetate/hexanes) to obtain the product as a viscous colorless oil (48.8 mg, 63.4 % yield): ^1^H NMR (500 MHz, d6-acetone) δ 10.6 (br s, 1), 8.64 (s, 1), 8.36 (d, *J* = 5.8 Hz, 1), 8.13 - 8.23 (m, 2), 7.90 - 8.05 (m, 2), 7.79 (t, *J* = 7.5 Hz, 1), 7.74 (d, *J* = 5.5 Hz, 1), 7.67 (t, *J* = 7.6 Hz, 1), 2.70 - 2.90 (m, 2 + H2O), 1.71 (quin, *J* = 7.6 Hz, 2), 1.35 - 1.50 (m, 2), 0.97 (t, *J* = 7.3 Hz, 3). Calculated for C19H20N3O (M+H)^+^ = 306.2; Observed 306.2.

A similar procedure was followed for the synthesis of compounds, utilizing POA and the appropriate amine, described below.

#### JSF-4301

^1^H NMR (500 MHz, CDCl3) δ 10.1 (d, *J* = 44.7 Hz, 1), 9.50 (s, 1), 8.80 (s, 1),

8.60 (s, 1), 8.39 (d, *J* = 8.3 Hz, 1), 8.36 (d, *J* = 4.6 Hz, 1), 7.77 (t, *J* = 7.8 Hz, 1), 7.10 (t,

*J* = 5.9 Hz, 1). Calculated for C10H9N4O (M+H)^+^ = 201.1; Observed 201.0.

#### JSF-4302

^1^H NMR (500 MHz, d6-dimethyl sulfoxide (DMSO)) δ 11.2 (s, 1), 9.31 (s, 1),

8.99 (d, *J* = 2.4 Hz, 1), 8.87 (s, 1), 8.41 (d, *J* = 5.7 Hz, 1), 8.06 (dd, *J* = 11.2, 8.6 Hz, 2),

7.90 – 7.79 (m, 2), 7.68 (t, *J* = 7.6 Hz, 1). Calculated for C14H11N4O (M+H)^+^ = 251.1; Observed 251.0.

#### JSF-4303

^1^H NMR (500 MHz, CDCl3) δ 10.1 (d, J = 45.4 Hz, 1), 9.52 (s, 1), 8.82 (s, 1), 8.62 (s, 1), 8.31 (d, J = 8.4 Hz, 1), 8.20 (s, 1), 7.61 (d, J = 8.4 Hz, 1), 2.35 (s, 3). Calculated for C11H11N4O (M+H)^+^ = 215.1; Observed 215.0.

#### JSF-4304

^1^H NMR (500 MHz, CDCl3) δ 10.1 (s, 1), 9.51 (s, 1), 8.82 (s, 1), 8.62 (s, 1), 8.26 (s, 1), 8.22 (d, J = 4.9 Hz, 1), 6.94 (d, J = 4.9 Hz, 1), 2.43 (s, 3). Calculated for C11H11N4O (M+H)^+^ = 215.1; Observed 215.0.

### Minimum inhibitory concentration determination

The MIC values of Mtb H37Rv, Mtb clinical isolates, and BCG were measured in 96-well microtiter plates using the alamarBlue™ assay as previously described (50). Briefly, strains were grown to mid-log phase were diluted (1:100) in Middlebrook 7H9 supplemented with 10% albumin, dextrose, sodium chloride (ADS) (Sigma) and 0.05% tyloxapol (Sigma). A 50 μL aliquot of this dilution was then added to each well that contained 11 2-fold serial dilutions of test compound. After 7 d of incubation at 37 °C, a mixture of alamarBlue™ (ThermoFisher Scientific) reagent and 20% Tween 80 (Sigma) was added to each well to evaluate bacterial cell viability. Plates were scanned after 24 h at 570 nm absorbance with a reference wavelength of 600 nm.

BD BACTEC PZA Media (pH 5.9) tubes were supplemented with 800 μL of BD BACTEC OADC and 100 μL of drug at the desired testing concentration. Mtb strains were grown to mid-log phase, diluted to 10^5^ CFU/mL, and 100 μL was added to each tube. For the drug-free control tubes, strains were diluted 1:100 in Middlebrook 7H9 and then 100 μL was added to each tube. An MIC99 value was defined as the first concentration needed for the culture to reach positivity after the 1:100 diluted control culture.

### Isolation of resistant mutants and their genetic characterization

Mid-log phase Mtb cultures (10^8^ CFU/mL) were plated on Middlebrook 7H10 agar containing drug at 4x, 8x, 16x, and 32x the MIC and incubated for 6 weeks at 37 °C. Isolated colonies were purified and grown in liquid culture and subjected to MIC determination. Genomic DNA was isolated from confirmed resistant mutants and subjected to whole-genome sequencing using Illumina NextSeq at 100x coverage. Variant calling was performed by DNAstar using the H37Rv sequence AL123456.3 as a reference. Sanger sequencing of *amiC* was conducted on an additional 12 DG160 and 2 DA60 mutants.

### Construction of overexpression strains

The *amiC* gene was amplified using polymerase chain reaction (PCR) from WT H37Rv using forward primer (GGAATCACTTCCATATGTCGCGCGTACACGCTTTCG) and reverse primer (GGTGGTGGTGAAGCTTCTCGGCGATATTTGGGGCG). The resulting PCR product was then cloned into pVV16-hsp60 between NdeI and HinDIII using In-Fusion® HD cloning kit (Takara Bio USA, Inc) and Stellar™ competent cells. The plasmids were prepared, and sequences were verified using Sanger sequencing. Once clones were confirmed, the plasmids were transformed into Mtb H37Rv using electroporation.

### AmiC purification

A Mtb H37Rv strain containing an overexpressing His-tagged AmiC was grown to mid-log phase (OD595 = 0.6 - 1.0) in Middlebrook 7H9 + 10% OADC + 0.2% glycerol + 0.05% Tween 80. The culture was then spun at 4,000 xg for 10 min, the supernatant was discarded, and the pellet was resuspended in 1x phosphate buffer saline. The sample was transferred to a bead beating tube and beat 3x at maximum speed for 30 s. Tubes were then centrifuged at 5,000 xg for 20 min at 4°C. Next, the supernatant was spun through a Spin-X 0.2 μM filter column (Costar) at 4,000 xg for 10 min. His-tagged AmiC was isolated from the protein sample using a TALON® metal affinity resin (Takara Bio USA, Inc) as previously described (51) and Sephadex 200 gel filtration.

### Intrabacterial drug metabolism assay

The IBDM assays were conducted as previously described (39, 40). Briefly, strains were grown to mid-log phase (OD595 = 0.6 – 1.0) in Middlebrook 7H9 + 10% ADS + 0.05% tyloxapol and at each timepoint, 10 mL of culture was centrifuged at 4,000 xg for 10 min at 4 °C. The supernatant was discarded, and the pellet was then quenched with pre-chilled acetonitrile (ACN):methanol (MeOH):water (2:2:1) on dry ice for 30 s. The remaining pellet solution was lysed via bead beating (6.5 m/s, 30 s, x6) and placed on ice for 2 min in between each cycle to avoid overheating. Bead beating tubes were then centrifuged at 6,000 xg for 10 min at 4 °C. The organic solvent was then centrifuged through 0.22 μm filter tubes at 4,000 xg for 15 min at 4 °C.

### Liquid chromatography tandem mass spectrometry cellular assays

Neat 1 mg/mL stocks of test compounds in DMSO, were serial diluted 1:1 in ACN:water to create standard curves and quality control (QC) spiking solutions. Standards and QCs were created by adding 10 µL of spiking solutions to 90 µL of blank cell lysate or blank media. Extraction was performed by adding 10 µL of cell lysate or media to 100 µL of MeOH:ACN (1:1) precipitation solvent containing 10 ng/mL of the internal standard Verapamil. Extracts were vortexed for 5 min and centrifuged at 4,000 xg for 5 min. A 75 µL aliquot of supernatant was transferred for LC-MS/MS analysis and diluted with 75 µL of Milli-Q deionized water.

LC-MS/MS analysis was performed on a Sciex Applied Biosystems Qtrap 6500+ triple-quadrupole mass spectrometer coupled to a Shimadzu Nexera X2 UHPLC system to quantify metabolites each in samples. Chromatography was performed on a Luna Omega Polar C18 column (2.1x100 mm; particle size, 3 µm) using a reverse phase gradient. Milli-Q deionized water with 0.1% formic acid was used for the aqueous mobile phase and 0.1% formic acid in ACN for the organic mobile phase. Multiple-reaction monitoring of parent/daughter transitions in electrospray positive-ionization mode was used to quantify the analytes. Sample analysis was accepted if the concentrations of the quality control samples were within 20% of the nominal concentration. Data processing was performed using Analyst software (version 1.6.2; Applied Biosystems Sciex).

LC-MS analysis of AmiC extracts was performed on a Q-Exactive high-resolution mass spectrometer (QE-HRMS) at 70000 mass resolution using an Ultimate 3000 UHPLC system for chromatographic separation. Chromatographic conditions were the same. Amidase products were verified by extracted ion chromatogram of predicted amidase products using a 5 parts per million mass accuracy.

## Acknowledgments

This work was supported in part by NIH grants R33AI11167 (D.A.), R21AI111647 (D.A.), U19AI109713 (D.A., J.S.F., V.D.) and U19AI142731 (D.A., J.S.F., V.D.), 1S10OD026890-01A1 (D.A.). We thank GSK and SRI for the subset of anti-tuberculars for screening. We thank Dr. Thomas Dick for providing the *M. bovis* BCG POA-resistant strain.

## Author Contributions

C.L., J.S.F., D.A, and P.K. conceived and designed experiments; R.J, D.Aw., A.S., S.S.D, and R.J.D synthesized compounds; C.L, K.T., R.S., T.R., C.G., P.S., T.K., R.R., X.W., P.B., Y.P., J.H., M.Z., and V.D. performed biological experiments; C.L, J.S.F., D.A. and P.K. wrote the manuscript. All authors discussed the results, commented, and contributed to sections of the manuscript.

## SUPPLEMENTARY MATERIAL

**Table S1:** DG160 MIC values in clinical isolates

| Strain |  |  | alarmarBlue MIC (µg/mL) |
| --- | --- | --- | --- |
| Name | Drug Resistance | amiC Mutation | DG160 |
| TDR1 | RIF, EMB | NA | 0.61 |
| TDR6 | RIF, INH, EMB | NA | 0.61 |
| TDR7 | RIF, INH, EMB, STR | NA | 0.31 |
| TDR19 | RIF, INH, EMB | P355fs | 1.2 |
| TDR21 | RIF, INH, EMB | NA | 0.31 |
| TDR26 | RIF, INH | NA | 0.15 |
| TDR29 | NA | NA | 0.15 |
| TDR31 | RIF, INH, EMB | NA | 0.15 |
| TDR32 | RIF, INH, EMB, STR | NA | 0.038 |
| TDR36 | RIF, INH, EMB | NA | 0.31 |
| TDR37 | INH, EMB | NA | 0.31 |
| TDR42 | INH, EMB | NA | 0.31 |
| TDR50 | RIF, INH, EMB | NA | 0.15 |
| TDR58 | RIF, INH, EMB | NA | 0.15 |
| TDR67 | RIF, INH, EMB | NA | 0.15 |
| TDR72 | RIF, INH, EMB | NA | 0.31 |
| TDR73 | RIF, INH, EMB | NA | 0.31 |
| TDR74 | RIF, INH, EMB | NA | 0.61 |
| TDR87 | RIF, EMB | NA | 0.31 |
| TDR91 | NA | H294fs | 4.9 |
| TDR92 | NA | Q399fs | 4.9 |
| TDR93 | INH, EMB | Q399fs | 4.9 |
| TDR101 | RIF, INH, EMB | NA | 0.61 |
| TDR104 | INH, EMB | NA | 0.15 |
| TDR105 | INH, EMB | NA | 0.15 |
| TDR106 | RIF, INH, EMB | NA | 0.31 |
| TDR109 | INH, EMB, STR | NA | 0.31 |
| TDR114 | RIF, INH, EMB | NA | 0.31 |
| TDR116 | RIF, INH, EMB | NA | 0.61 |
| TDR120 | EMB | Q399fs | 4.9 |
| TDR123 | INH, EMB, STR | NA | 0.15 |
| TDR124 | RIF, INH | NA | 0.61 |
| TDR126 | EMB | NA | 0.61 |
| TDR128 | EMB | NA | 0.31 |
| TDR132 | INH, EMB | NA | 0.31 |
| TDR133 | RIF, EMB | NA | 0.15 |
| TDR138 | EMB | NA | 0.15 |
| TDR143 | EMB | NA | 0.15 |
| TDR146 | EMB | NA | 0.15 |
| TDR150 | RIF, INH, EMB | NA | 0.61 |
| TDR156 | NA | NA | 0.61 |
| TDR162 | RIF, INH, EMB | NA | 0.15 |
| TDR173 | EMB | NA | 0.61 |
| TDR191 | RIF, EMB | NA | 0.31 |
| TDR192 | RIF, INH, EMB | NA | 0.31 |
| TDR195 | INH, EMB | NA | 0.31 |
| TDR196 | RIF, INH, EMB, STR | NA | 0.15 |
| TDR216 | RIF, INH, EMB | NA | 0.31 |

**Table S2:**
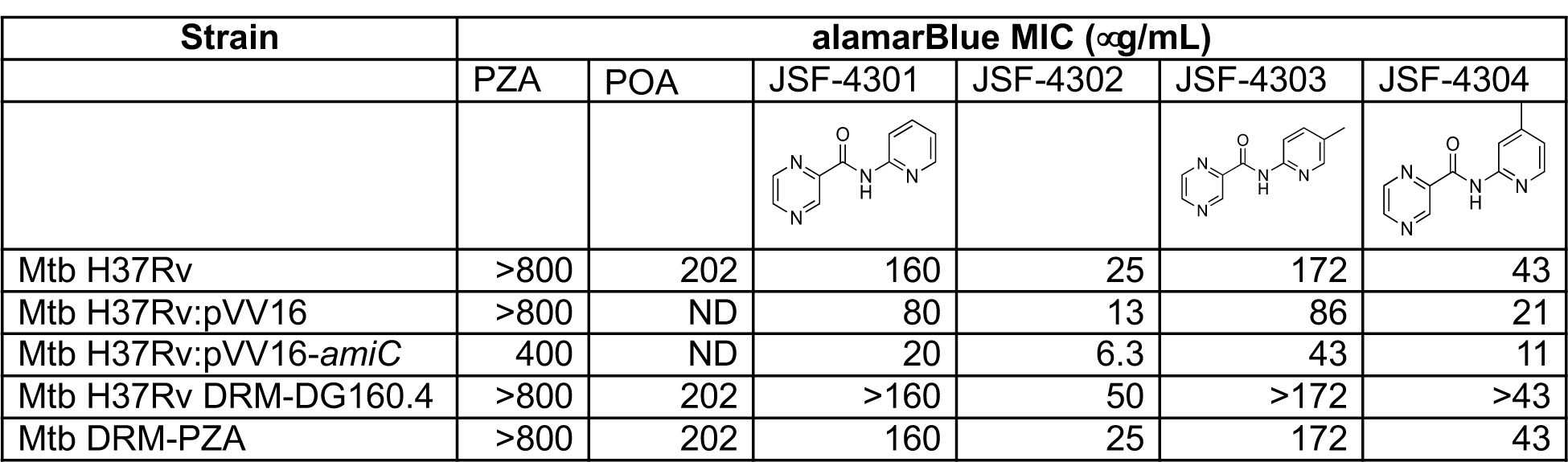
MIC values for DG160-POA compounds at neutral pH

| Strain | alamarBlue MIC (μg/mL) | | | | | |
| --- | --- | --- | --- | --- | --- | --- |
|  | PZA | POA | JSF-4301 | JSF-4302 | JSF-4303 | JSF-4304 |
| Mtb H37Rv | >800 | 202 | 160 | 25 | 172 | 43 |
| Mtb H37Rv:pVV16 | >800 | ND | 80 | 13 | 86 | 21 |
| Mtb H37Rv:pVV16-*amiC* | 400 | ND | 20 | 6.3 | 43 | 11 |
| Mtb H37Rv DRM-DG160.4 | >800 | 202 | >160 | 50 | >172 | >43 |
| Mtb DRM-PZA | >800 | 202 | 160 | 25 | 172 | 43 |

**Table S3:**
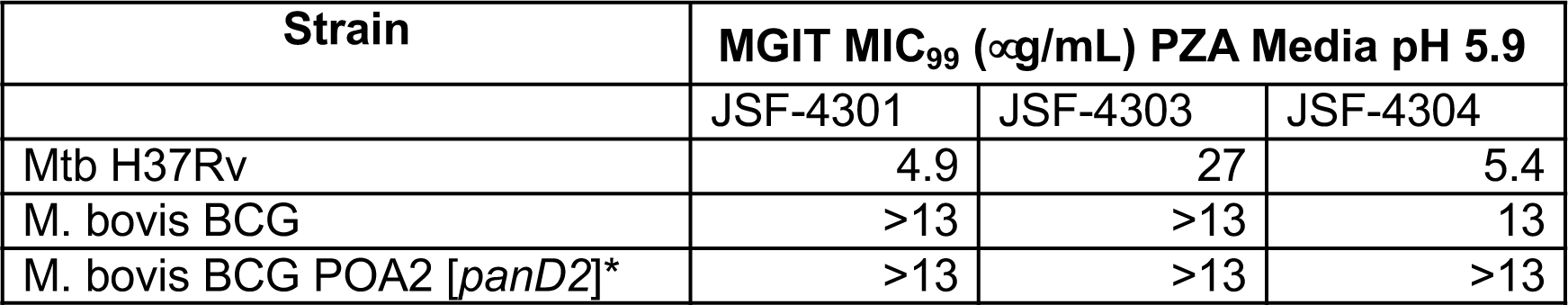
MIC_99_ values for DG160-POA compounds at acidic pH

| Strain | MGIT MIC<sub>99</sub> (μg/mL) PZA Media pH 5.9 | | |
| --- | --- | --- | --- |
|  | JSF-4301 | JSF-4303 | JSF-4304 |
| Mtb H37Rv | 4.9 | 27 | 5.4 |
| M. bovis BCG | >13 | >13 | 13 |
| M. bovis BCG POA2 [*panD2*]* | >13 | >13 | >13 |

**Fig. S1:**
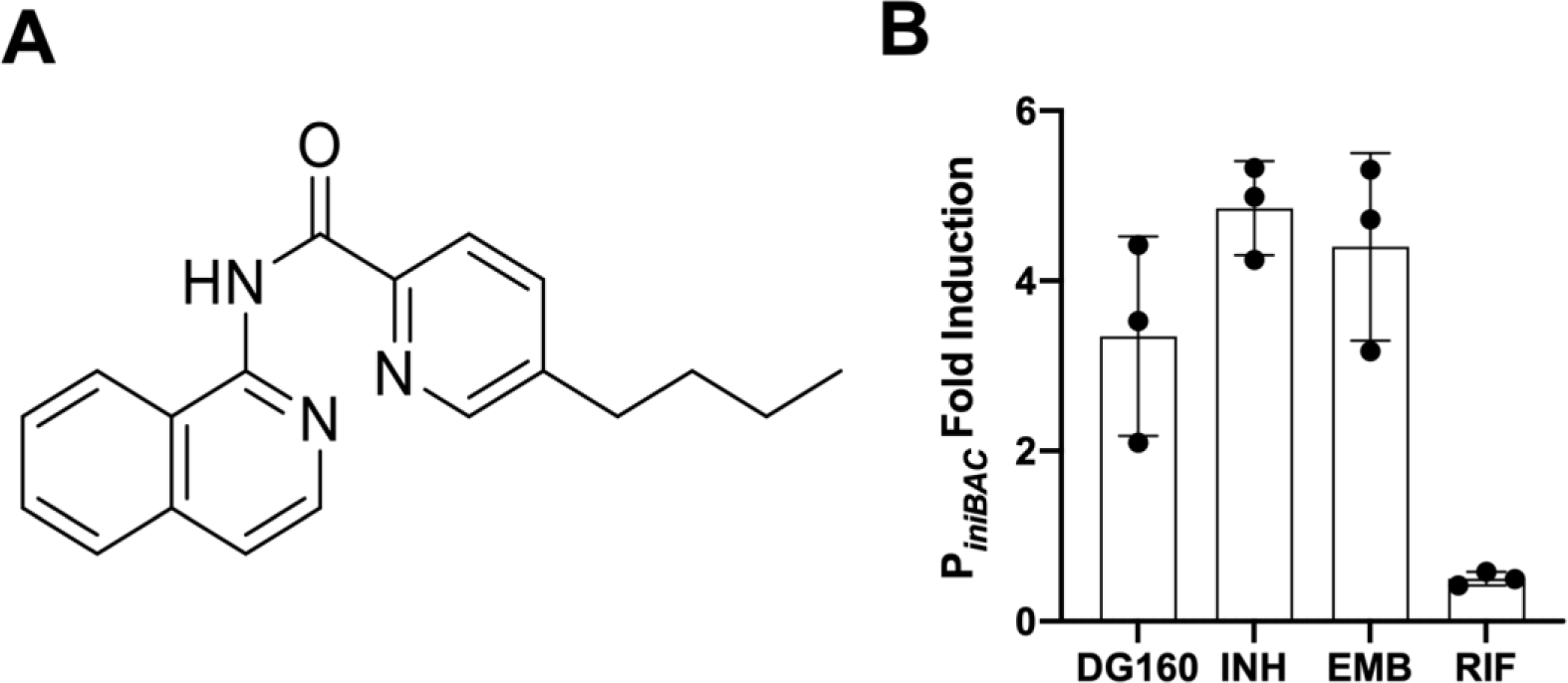
DG160 is an inducer of the *iniBAC* operon. **A)** Structure of DG160, **B)** *M. bovis* BCG P*iniBAC* screen identified DG160 as an inducer of the *iniBAC* operon 24 h post treatment. INH and EMB included as positive controls. RIF included as a negative control. Columns and error bars represent the mean and standard deviation, respectively, from n = 3 wells.

**Fig. S2:**
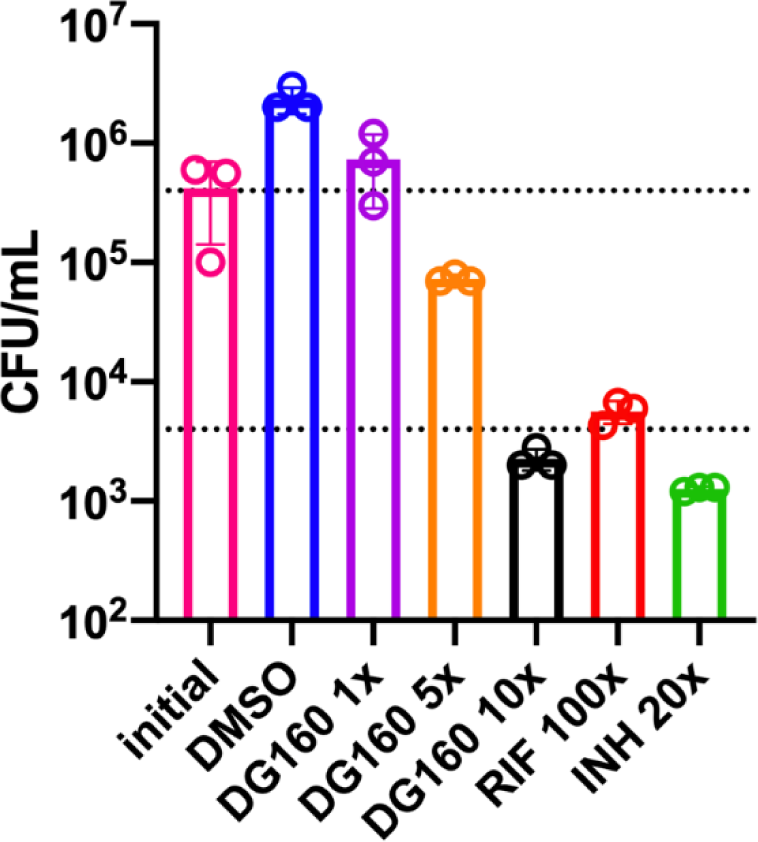
DG160 is bactericidal. Reduction of 2 log_10_ CFU/mL (represented by dotted lines) 4 d post treatment with 10x MIC DG160. Data points and error bars represent the mean and standard deviation, respectively. At all concentrations n = 3.

**Fig. S3:**
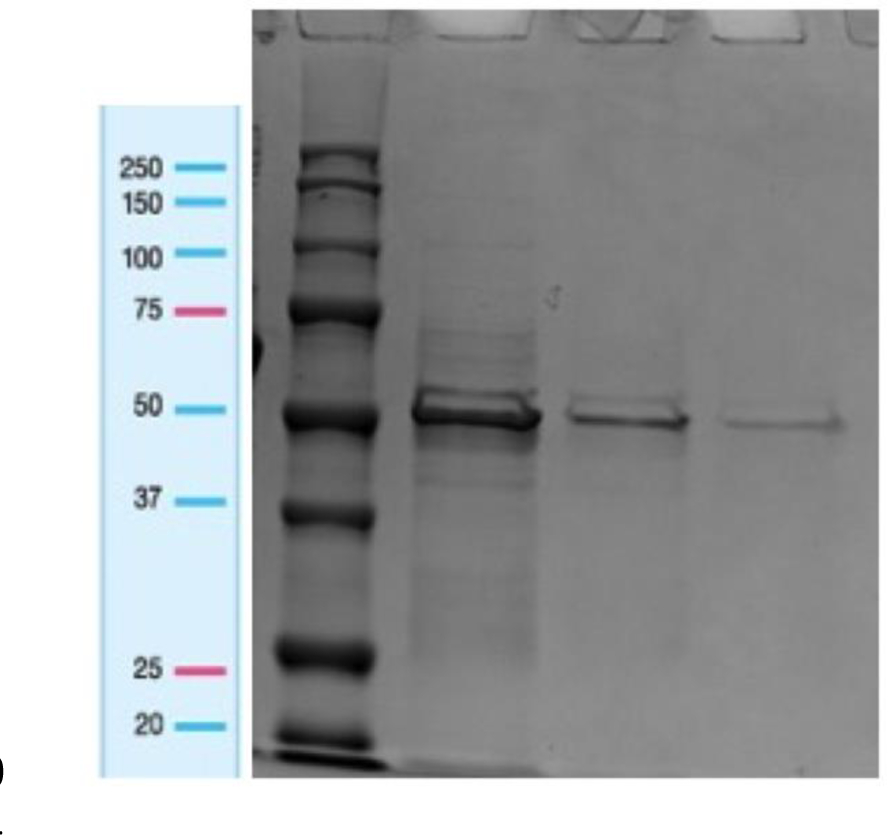
AmiC isolated from Mtb H37Rv:pVV16-*amiC*. SDS-PAGE of isolated His-tagged AmiC. Dark band at ∼50 kD corresponds with AmiC (MW of 50.92 kD).

**Fig. S4:**
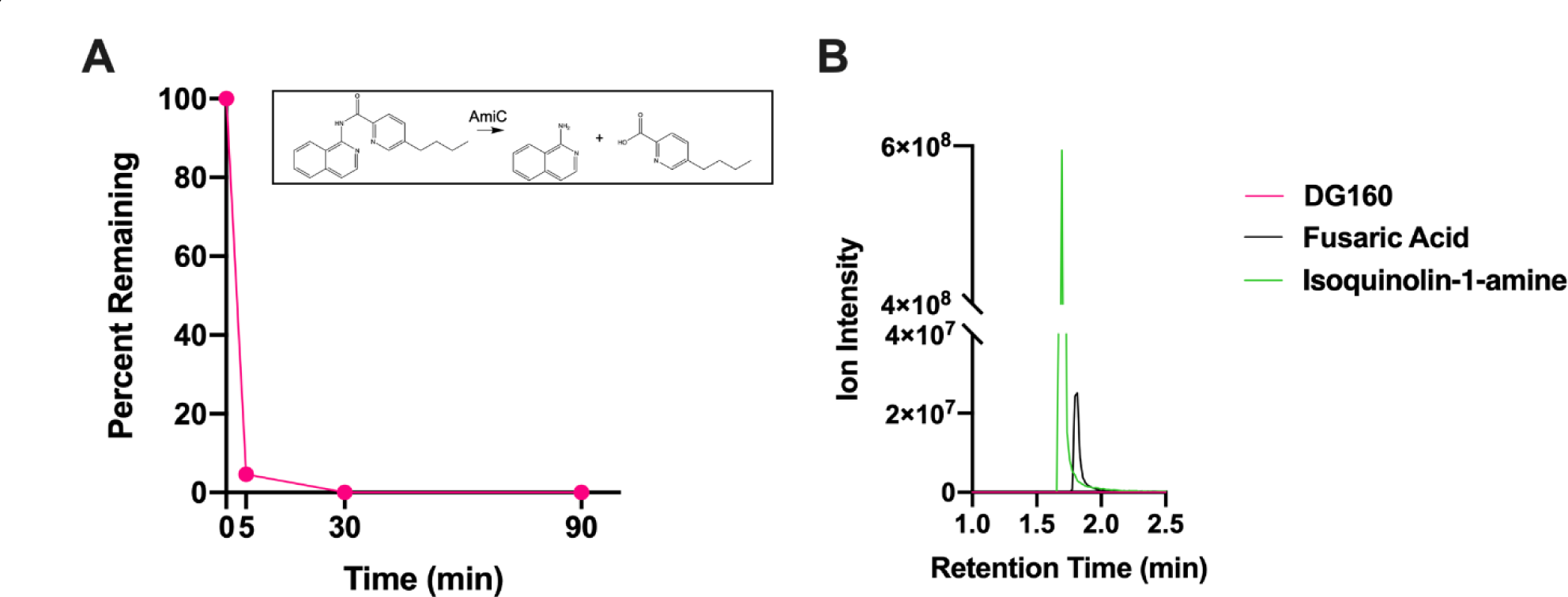
DG160 is metabolized at the amide bond by isolated AmiC. A) Percent DG160 remaining over time, **B)** EIC of DG160, fusaric acid, and isoquinolin-1-amine at 90 min incubation with AmiC. Graphs are representative of n = 3 experiments.

**Fig. S5:**
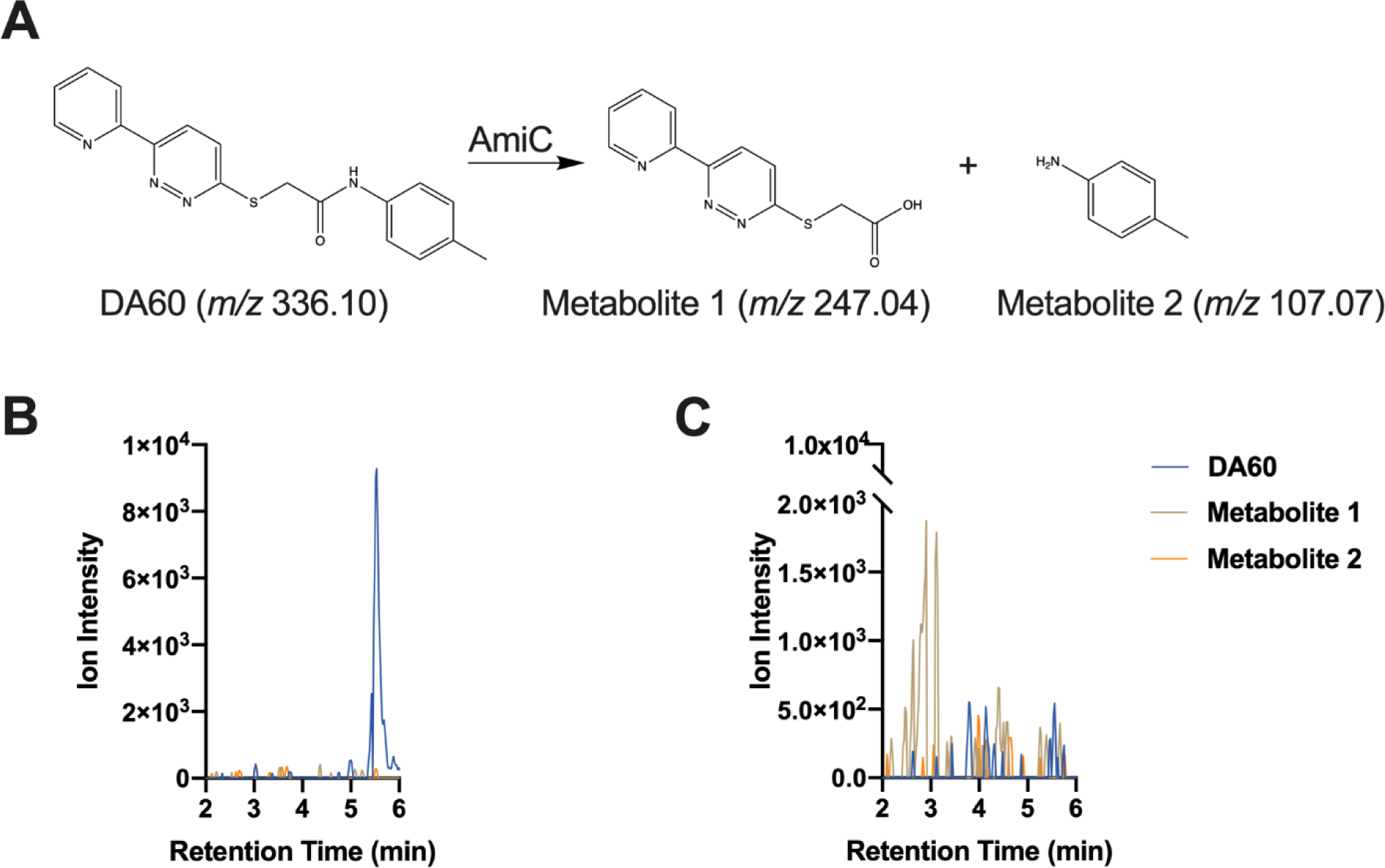
DA60 is metabolized at the amide bond by isolated AmiC. A) Schematic of amide bond hydrolysis and subsequent generation of Metabolite 1 and Metabolite 2, **B)** EIC of DA60, Metabolite 1, and Metabolite 2 after 2 h incubation with heat killed AmiC, **C)** EIC of DA60, Metabolite 1, and Metabolite 2 after 2 h incubation with AmiC.

**Fig. S6:**
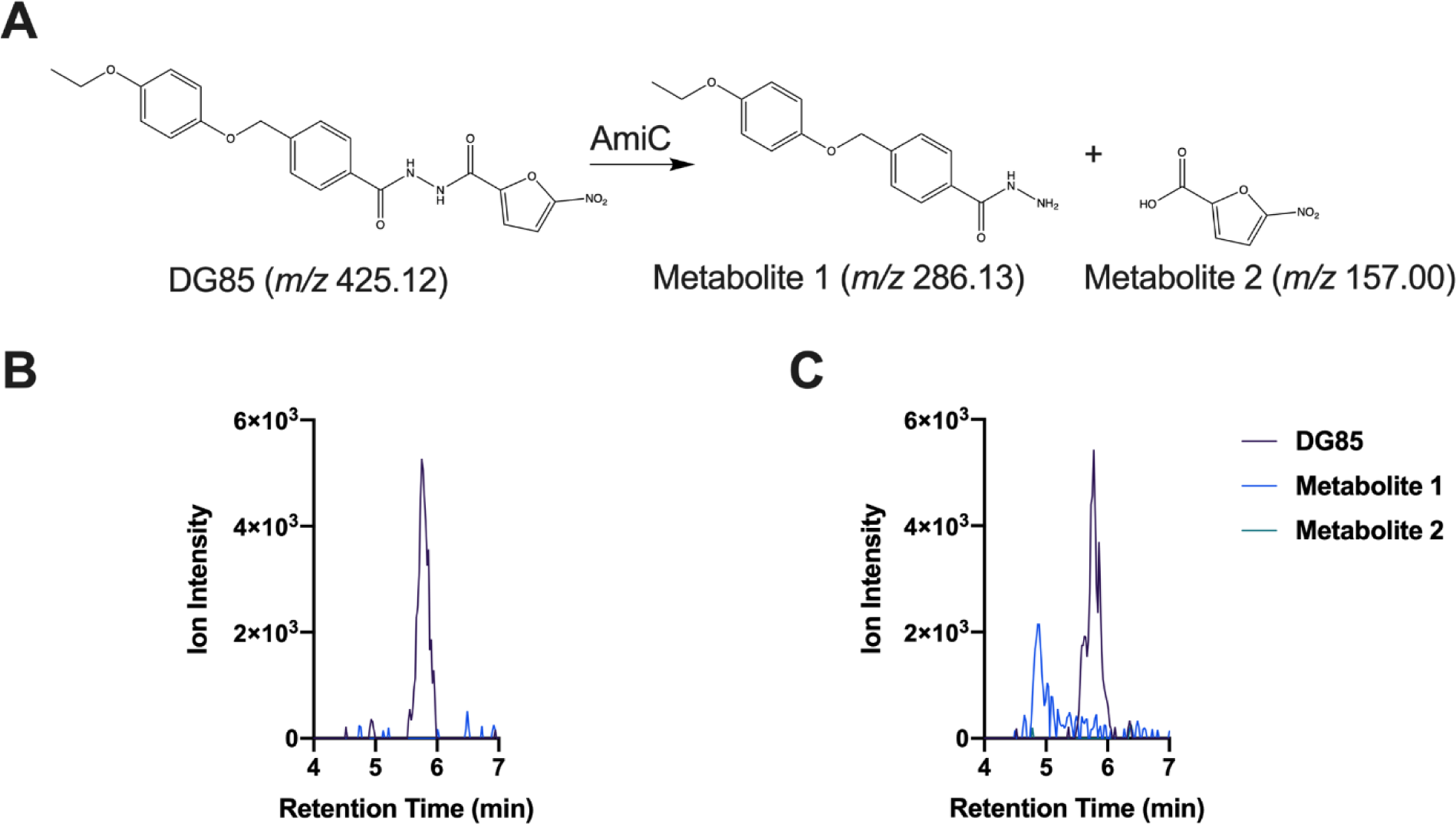
DG85 is metabolized at the amide bond by isolated AmiC. **A)** Schematic of amide bond hydrolysis and subsequent generation of Metabolite 1 and Metabolite 2, **B)** EIC of DG85, Metabolite 1, and Metabolite 2 after 2 h incubation with heat killed AmiC, **C)** EIC of DG85, Metabolite 1, and Metabolite 2 after 2 h incubation with AmiC.

**Fig. S7:**
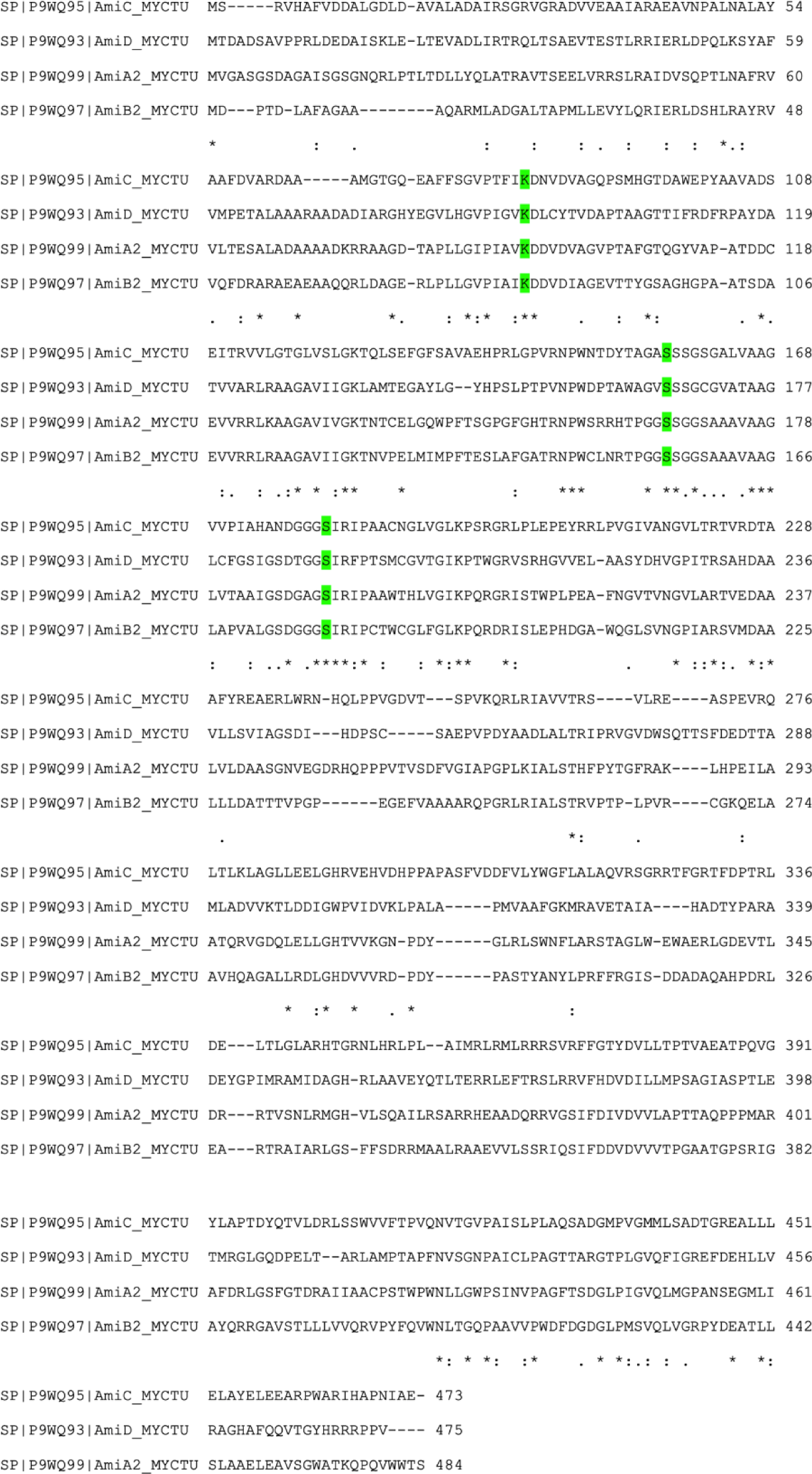
AmiC homologs based on a conserved KSS amidase family signature. Clustal Omega multiple sequence analysis showing conserved KSS residues in putative homologs in Mtb.

**Fig. S8:**
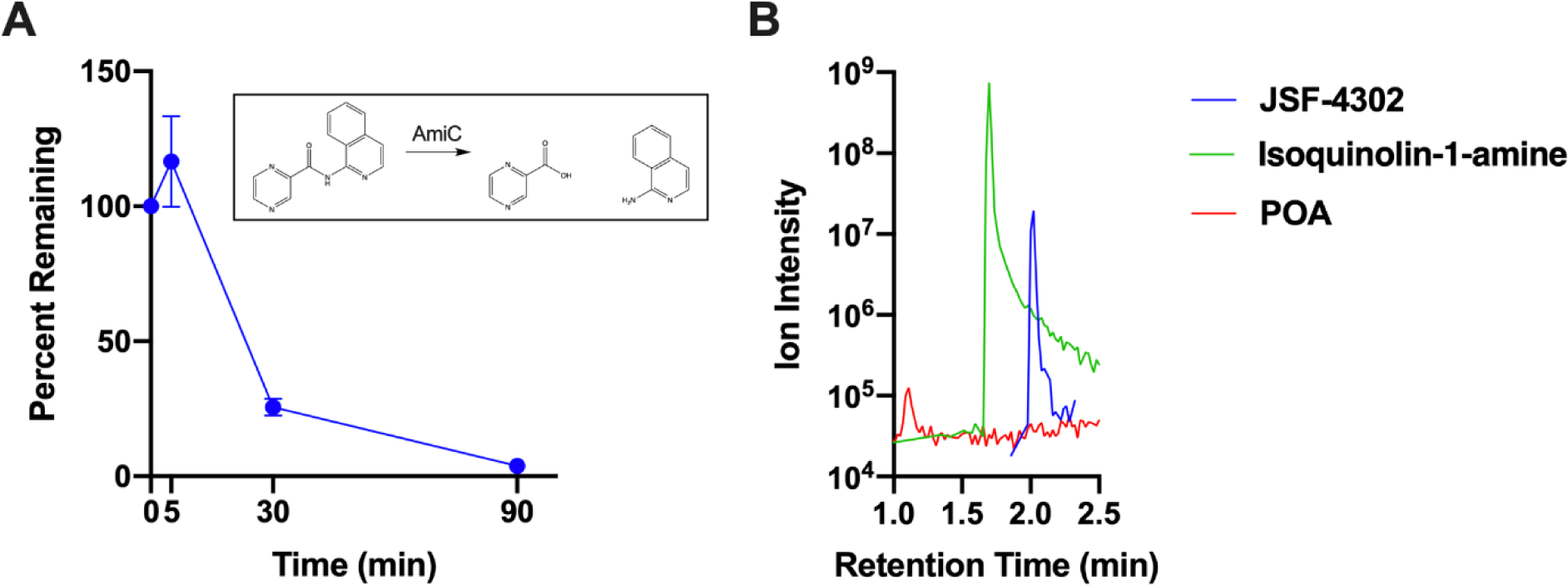
JSF-4302 is metabolized at the amide bond by isolated AmiC. A) Percent JSF-4302 remaining over time, **B)** EIC of JSF-4302, JSF-4136, and POA after 90 min incubation with AmiC. Graphs are representative of n = 3 experiments.

